# Insights into the mysterious genetic variation profile of *tprK* in *Treponema pallidum* under the development of natural human syphilis infection

**DOI:** 10.1101/536573

**Authors:** Dan Liu, Man-Li Tong, Yong Lin, Li-Li Liu, Li-Rong Lin, Tian-Ci Yang

## Abstract

Although the variations of the *tprK* gene in *Treponema pallidum* were considered to play a critical role in the pathogenesis of syphilis, how actual variable characteristics of *tprK* in the course of natural human infection enabling the pathogen’s survive has thus far remained unclear. Here, we performed NGS to investigate *tprK* of *T. pallidum* directly from primary and secondary syphilis samples. Compared with diversity in *tprK* of the strains from primary syphilis samples, there were more mixture variants found within seven V regions of the *tprK* gene among the strains from secondary syphilis samples, and the frequencies of predominant sequences within V regions of *tprK* were generally decreased (less than 80%) with the proportion of minor variants in 10-60% increasing. Noteworthy, the variations within V regions of *tprK* always obeyed a strict 3 bp changing pattern. And *tprK* in the strains from the two-stage samples kept some stable amino acid sequences within V regions. Particularly, the amino acid sequences IASDGGAIKH and IASEDGSAGNLKH in V1 not only presented a high proportion of inter-population sharing, but also presented a relatively high frequency (above 80%) in the populations. Besides, *tprK* always demonstrated remarkable variability in V6 at both the intra- and inter-strain levels regardless of the course. These findings unveiled that the different profile of *tprK in T. pallidum* directly from primary and secondary syphilis samples, indicating that throughout the development of syphilis *T. pallidum* constantly varies its domain *tprK* gene to obtain the best adaptation to the host. While this changing was always subjected a strict gene conversion mechanism to keep an abnormal TprK. The highly stable peptides found in V1 would probably be promising potential vaccine components. And the highly heterogenetic regions (e.g. V6) could provide insight into the mysterious role of *tprK* in immune evasion.

**Author summary:** Although the variations of the *tprK* gene in *Treponema pallidum* were considered to play a critical role in the pathogenesis of syphilis, how actual variable characteristics of *tprK* in the course of natural human infection enabling the pathogen’s survive has thus far remained unclear. Here, we performed next-generation sequencing, a more sensitive and reliable approach, to investigate *tprK* of *Treponema pallidum* directly from primary and secondary syphilis patients, revealing that the profile of *tprK* in *T. pallidum* from the two-stage samples was different. Within the strains from secondary syphilis patients, more mixture variants within seven V regions of *tprK* were found, the frequencies of their predominant sequences were generally decreased with the proportion of minor variants in 10-60% was increased. And the variations within V regions of *tprK* always obeyed a strict 3 bp changing pattern. Noteworthy, the amino acid sequences IASDGGAIKH and IASEDGSAGNLKH in V1 presented a high proportion of inter-population sharing and presented a relatively high frequency in the populations. And V6 region always demonstrated remarkable variability at intra- and inter-patient levels regardless of the course. These findings provide insights into the mysterious role of TprK in immune evasion and for further exploring the potential vaccine components.

## Introduction

The natural history of syphilis is one of a complex chronic disease caused by the infection of *Treponema pallidum* subsp. *pallidum*. The disease has a series of highly distinct clinical stage [1], which usually includes the localized chancre primary stage, the disseminated secondary stage, and the late tertiary stage in untreated individuals [2]. This characteristic pattern of successive episodes is reminiscent of antigenic variation during pathogen infection that accounts for these repeated cycles of pathology [3, 4]. Studies have indicated that antigenic variation in outer membrane antigens is a hallmark of many chronic multistage infectious diseases [5–7].

Previous investigations of *tprK*, from a 12-member paralogue of the *T. pallidum* repeat (*tpr*) gene family, have revealed that *tprK* is highly heterogeneous at both the inter- and intra-strain levels, with sequence diversity in seven discrete variable regions (V1–V7) that are separated by conserved sequences [8–10]. Although a surface location for TprK is still controversial [11–13], many studies hypothesize that antigenic variation of TprK would facilitate *T. pallidum’s* ability to escape immune clearance, thus permitting the pathogen to persist in the host and remarkable results using rabbit models support this hypothesis [14–16]. In this regard, Reid *et al*. [17] explored the role of *tprK* in the development of secondary syphilis in a rabbit model based on a clone-based Sanger approach, demonstrating that the rampant variants of TprK were instrumental in the development of later stages of syphilis. However, the authors inevitably encountered a problem in that the inoculum did not maintain an exactly identical *tprK* clone as required. Additionally, important information about the variants, especially those with low-level diversity, would be lost if only the clone-based Sanger approach was used. Therefore, understanding of the variations of *tprK* that facilitate the development of syphilis is not complete. In our previous study [18], we employed next-generation sequencing (NGS) to explore the *tprK* gene directly from primary syphilis samples, demonstrating that the profile of *tprK* in primary syphilis patients generally contains a high proportion sequence (frequency above 80%) and many low-frequency minor variants (frequency below 20%) within each region. Only some sequences had frequencies between 20% and 80%. This causes us question whether this characteristic distribution of variants in *tprK* changes with the development of disease and whether *tprK* keep some relatively stable components in these rampant variations.

In the present study, we sought to perform a comprehensive investigation of characteristic variations of *tprK* in *T. pallidum* directly from syphilis patients with primary and secondary disease by employing NGS, thus revealing extensive information on the association of genetic variations of *tprK* in *T. pallidum* with disease progression, providing important insights into the immune evasion and persistence of this pathogen or potential vaccine component for human immunology study.

## Results

### 1. NGS of *tprK* directly from primary and secondary syphilis patients

The samples (n=28) were collected at Zhongshan Hospital, Xiamen University. Of the 28 samples, 14 samples (P1~14) were from patients diagnosed with primary syphilis, and 14 samples (S1~14) were from patients with secondary syphilis. The clinical information for all 28 patients is shown in Table 1. The qPCR data of target gene *tp0574* showed that the amount of treponemal DNA in each clinical sample was sufficient for amplification of the full length *tprK*. Based on the sequencing data of *tp0136*, most strains belonged to the SS-14-like group, and only five belonged to the Nichols-like group. The median sequencing depth of the *tprK* segment samples ranged from 9810.91 to 56676.38, and the coverage ranged from 99.34% to 99.61% (S1 Table).

**Table 1.** Clinical characteristic of the study participants

| Characteristic |  | Clinical stage |  |
| --- | --- | --- | --- |
|  |  | Primary (n=14) | Secondary (n=14) |
| Gender |  |  |  |
| Male | n (%) | 13 (92.9) | 10 (71.4) |
| Female | n (%) | 1 (7.1) | 4 (40.0) |
| Age (Mean±SD) |  | 53.9±13.7 | 39.7±15.7 |
| Serum RPR titer, Median (IQR) |  | 1:16 (1:4-1:32) | 1:32 (1:16-1:128) |
| Serum TPPA |  | Positive | Positive |
| Dark field microscopy |  | Positive | Positive |
| Clinical manifestations |  |  |  |
| Chancre |  | 13 | 0 |
| Lymphadenopathy |  | 1 | 0 |
| Condyloma |  | 0 | 11 |
| Erythema |  | 0 | 3 |

| Genetic group by *tp0136* |  |  |
| --- | --- | --- |
| Nichols-like group | 2 | 3 |
| SS14-like group | 12 | 11 |

Abbreviations: RPR, reactive plasma reagin; TPPA, *T. pallidum* particle agglutination; IQR, interquartile range;

### 2. The characteristic profile of *tprK* in *T. pallidum* from the samples of the two clinical stages

Using the strategy, distinct nucleotide sequences within seven V regions of the *tprK* gene were captured from each sample, and 491 sequences were obtained in 14 secondary syphilis samples, which was more than that captured in primary syphilis samples (335 in total) (S2 Table). The trends in the number of the total different sequences distributed in the seven V regions of the *tprK* gene were roughly the same between the two-stage samples; the highest and lowest numbers of different sequences were both found in V6 and V1 in two-stage samples, respectively (Fig 1). When the frequencies of distinct sequences within each V region in each strain were calculated, they generally contained a predominant sequence within the regions across all the samples (Fig 2). However, compared to the frequency of the predominant sequence in primary syphilis samples, the frequencies of the sequence within the variable regions of *tprK* was generally decreased in secondary syphilis samples, especially in V7, where the number of sequences with a frequency lower than 60% was increased (8/14 vs 1/14). Notably, the frequency of predominate sequences in V1 of all 28 samples remained almost higher than 80%.

**Figure 1.**
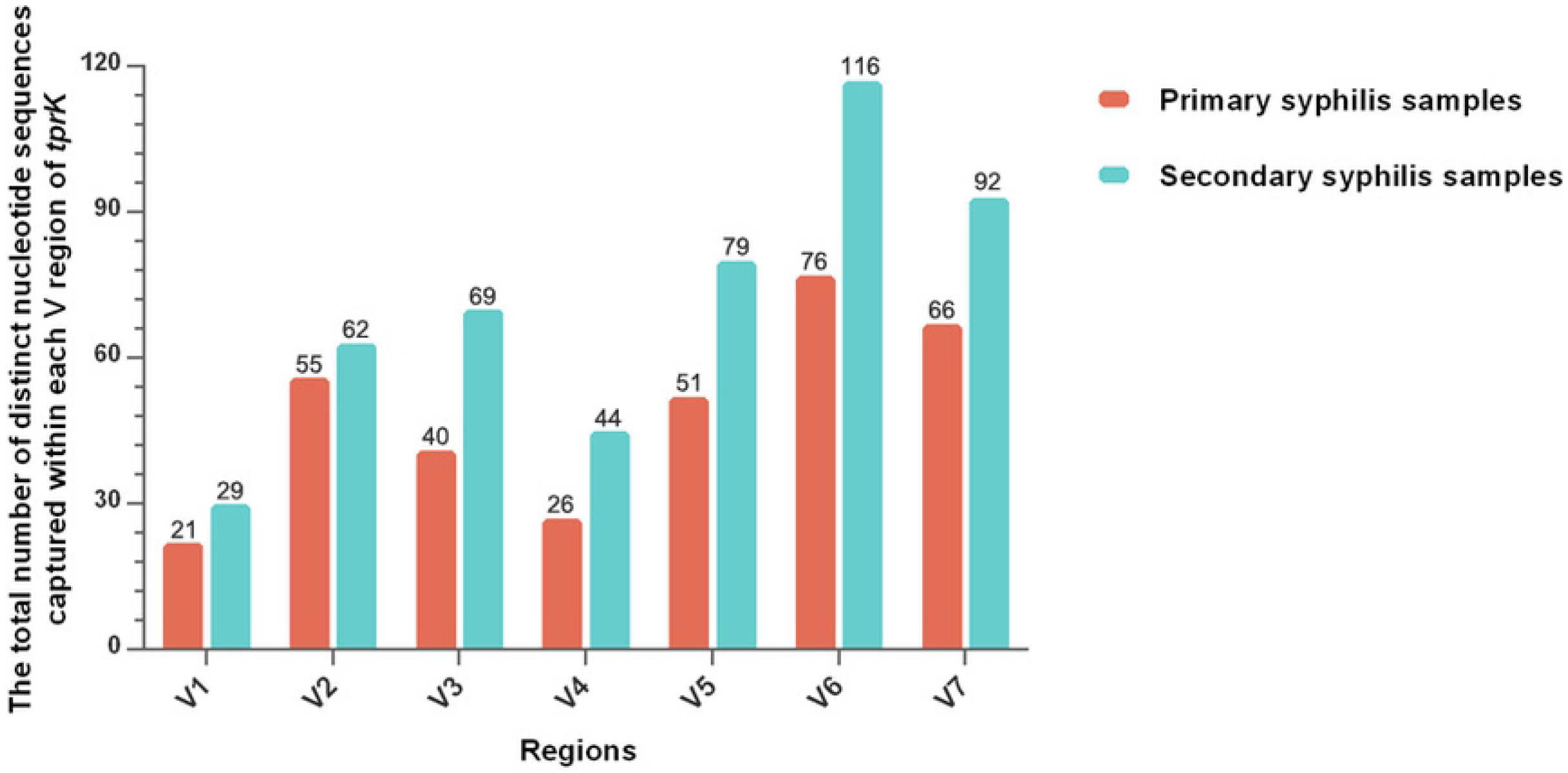
The number of distinct nucleotide sequences within each V region of each strain across the two-stage samples.

**Figure 2.**
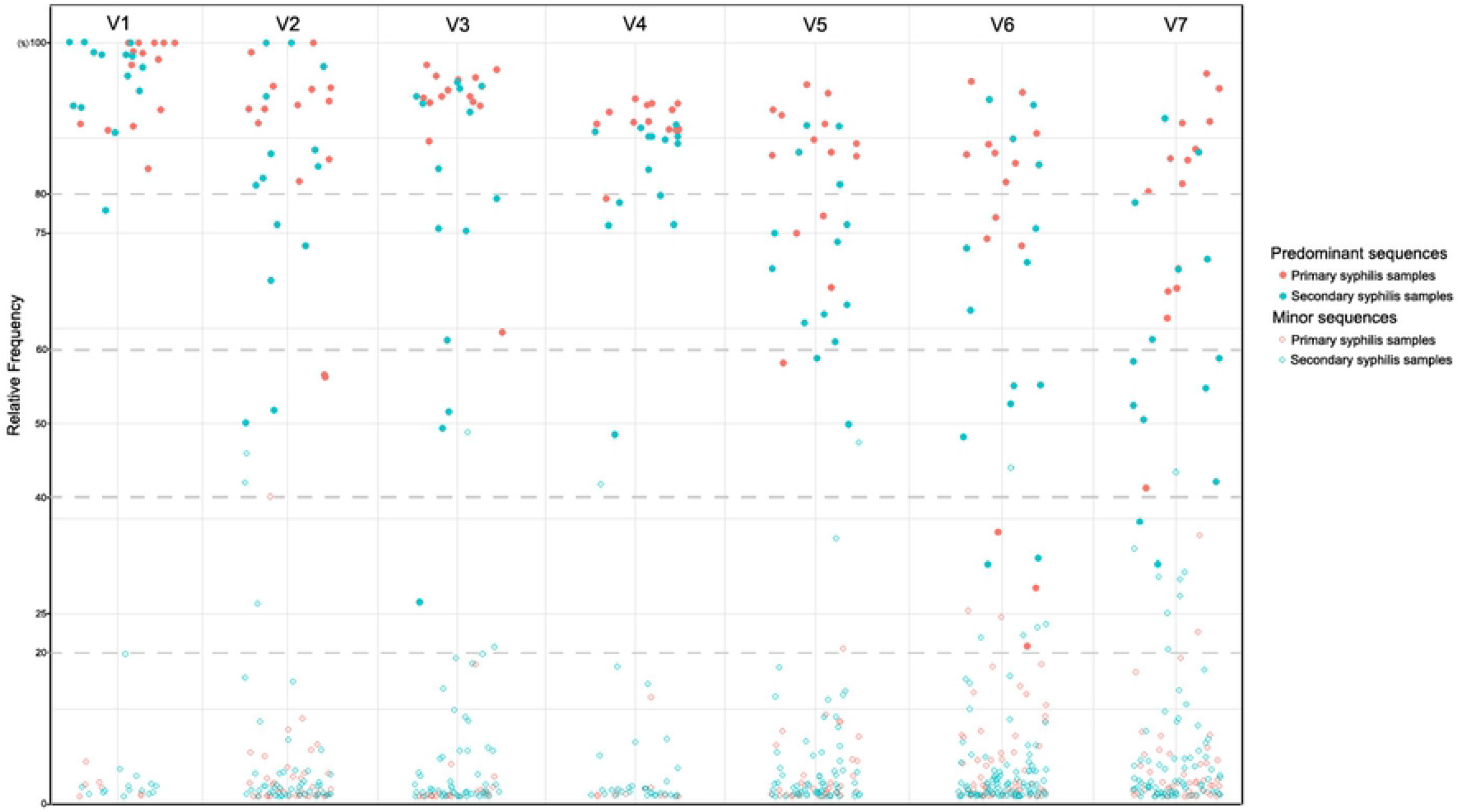
The proportion distribution of captured distinct nucleotide sequences within each V region of *tprK* from each sample. Each circle indicates the relative frequency of distinct sequences within each V region of *tprK*. The solid circle represents the predominant sequence and the hollow circles represent the minor variants in a particular population. The two colors specify the samples from the two clinical stages.

We applied three thresholds (1-5%, 5-10% and 10-60%) to investigate the distribution of minor variants. As Fig 3 shows, most of the minor variants were concentrated in the 1-5% range in both groups. However, the proportions in the other two ranges (5-10% and 10-60% in secondary syphilis samples) were reversed relative to the distribution pattern in primary syphilis samples; that is, the minor variants distributed in 10-60% unexpectedly increased in secondary syphilis samples; as a result, the lowest proportion range was replaced with 5-10% (37/393). Also, the proportion of minor variants distributed above 20% correspondingly increased, with 21/393 in the secondary syphilis samples relative to 6/237 in the primary syphilis samples.

**Figure 3.**
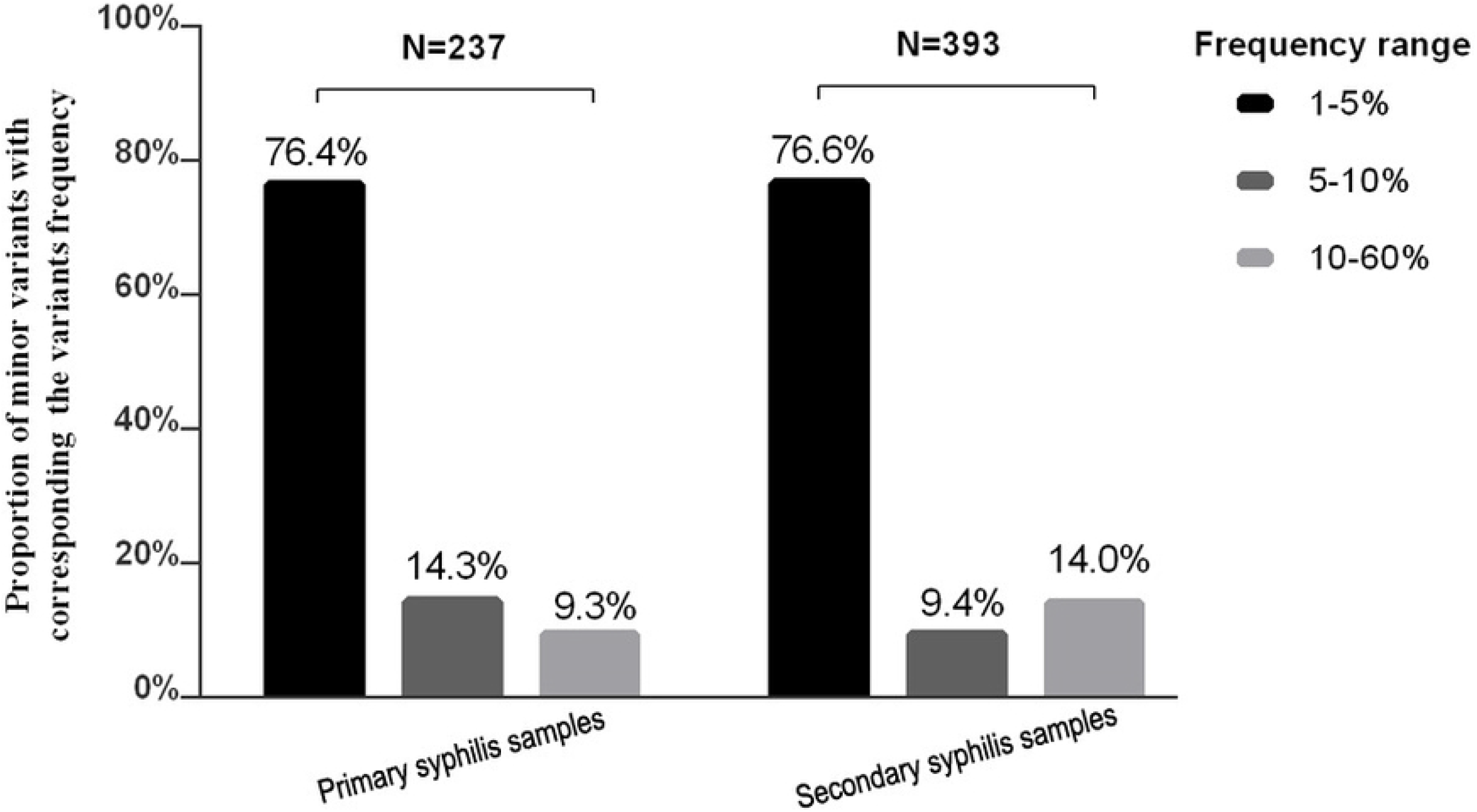
The total minor variants from primary and secondary syphilis samples distributed among the three thresholds (1-5%, 5-10% and 10-60%).

Then, the length of all captured sequences within the regions was analyzed, demonstrating that the change in length could be characterized as a 3 bp or multiple 3 bp pattern in either primary or secondary syphilis samples (Table 2). Compared to the forms of length within the regions in primary syphilis samples, V3 and V5 kept the same forms, while other regions in secondary syphilis samples had new lengths (V1, V4, V6 and V7) or both/or some lengths disappeared (V2 and V7). Particularly, the lengths of V5 in both different stage samples were found only in two lengths, 84 bp and 90 bp.

**Table 2.**
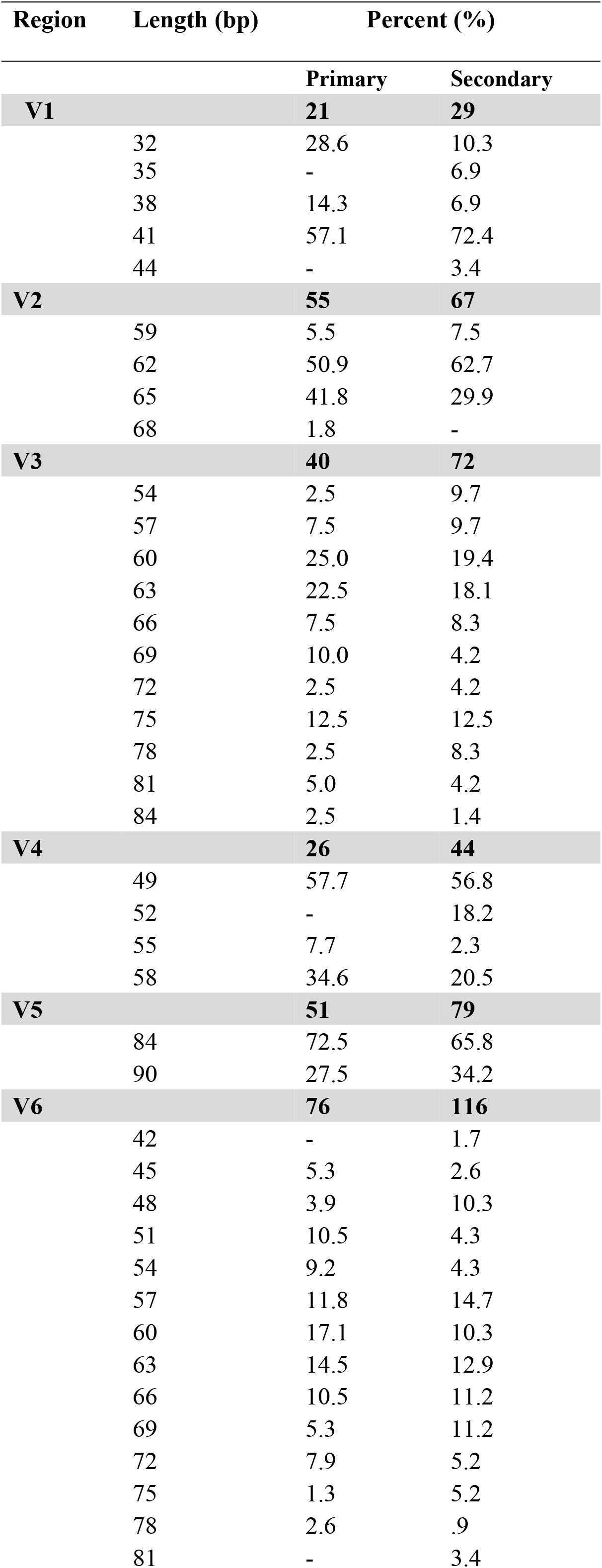
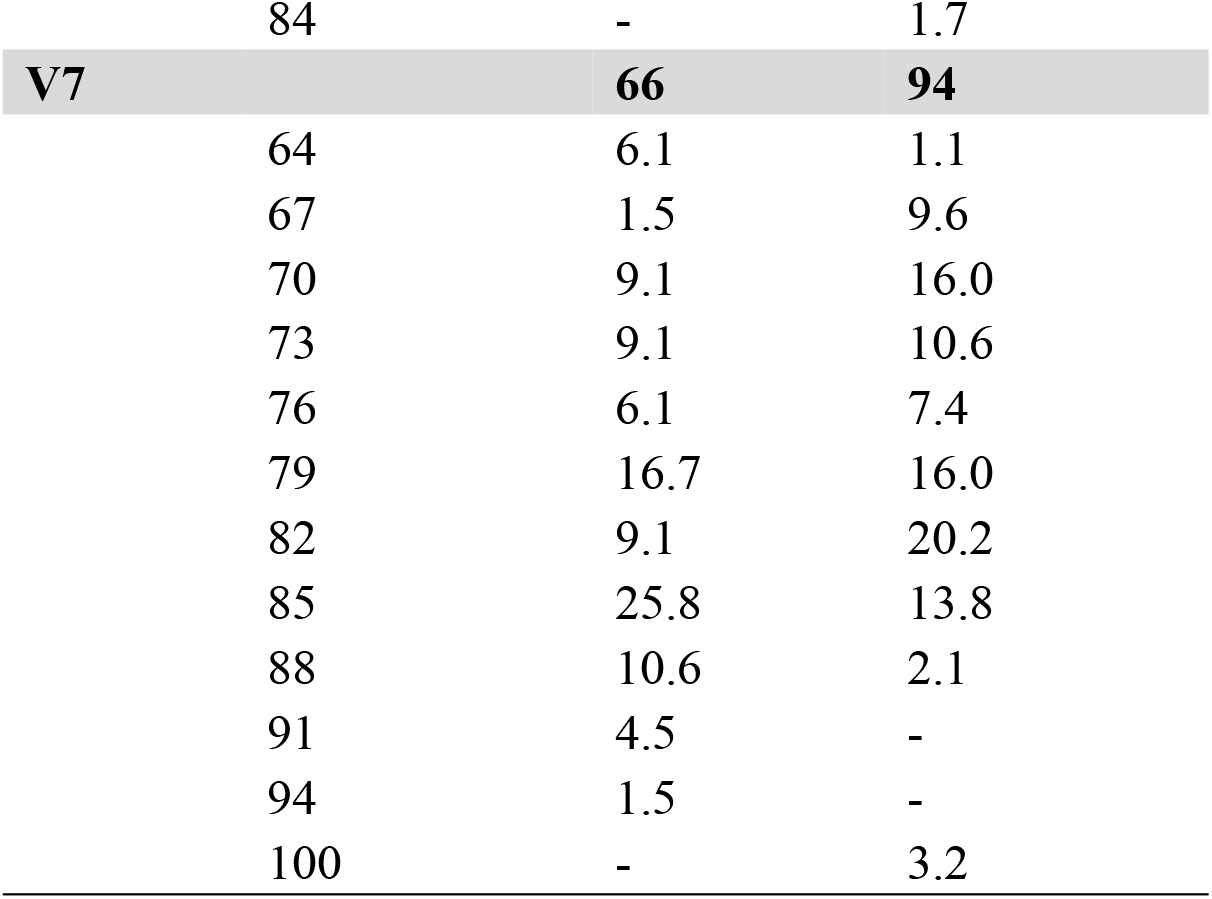
The length of distinct nucleotide sequences within seven V regions in *tprK* between primary and secondary syphilis samples

### 3. The amino acid sequence within each V region of the *tprK* gene

The captured nucleotide sequences within the seven V regions from each sample were translated into amino acid sequences *in silico*. There was no sequence yielding a *tprK* frame shift or premature termination, and synonymous sequences were rare and found only in V2 and V5 (S3 Table). Among the populations from the two-stage samples, a parallel scenario of sequence diversity in each V region was found. Altogether, V1, V2 and V4 had strong parallel sequence ability, and V6 was the region with the least consistent sequence diversity (Table 3).

**Table 3.** The interpopulation redundancy in amino acid sequences within seven V regions of *tprK* among primary and secondary syphilis samples

| V regions | Primary syphilis |  |  | Secondary syphilis |  |  |
| --- | --- | --- | --- | --- | --- | --- |
|  | Total distinct nucleotide sequence in single strain from 14 samples | No. case of synonymous mutations | Total type of sequences in 14 samples (%) | Total distinct nucleotide sequence in single strain from 14 samples | No. case of synonymous mutations | Total types of sequences in 14 samples (%) |
| V1 | 21 | 0 | 11 (52.4) | 29 | 0 | 14 (48.3) |
| V2 | 55 | 6 | 25 (45.5) | 62 | 8 | 23 (37.1) |
| V3 | 40 | 0 | 34 (85.0) | 69 | 0 | 54 (78.3) |
| V4 | 26 | 0 | 9 (34.6) | 44 | 0 | 14 (31.8) |
| V5 | 51 | 2 | 31 (60.8) | 79 | 4 | 46 (58.2) |
| V6 | 76 | 0 | 71 (93.4) | 116 | 0 | 110 (94.8) |
| V7 | 66 | 0 | 55 (83.3) | 92 | 0 | 66 (73.9) |

When all the exclusive types of amino acid sequences within the regions from the two-stage samples were aligned, some sequence types were continually found across primary and secondary syphilis samples (S4 Table). As described above, V1, V2 and V4 presented a strong inter-population shared sequence capacity, regardless of what stage the samples came from. Interestingly, the sequences in these three regions, especially in V2, showed a high proportion of identity across the samples from both groups (Fig 4a). Moreover, there contained predominant sequences of some populations among the observed identical types of sequences (Fig 4b). And some identical predominant amino acid sequences between the two-stage populations were the ones that had a high proportion of inter-population sharing. For example, the sequences DVGHKKENAANVNGTVGA and DVGRKKDGAQGTVGA in V4 were the two most stable sequences found among primary and secondary syphilis samples, respectively. The same phenomenon was observed in the sequences IASDGGAIKH and IASEDGSAGNLKH of V1. Moreover, the frequency of these sequences in V1 was above 80%.

**Figure 4.**
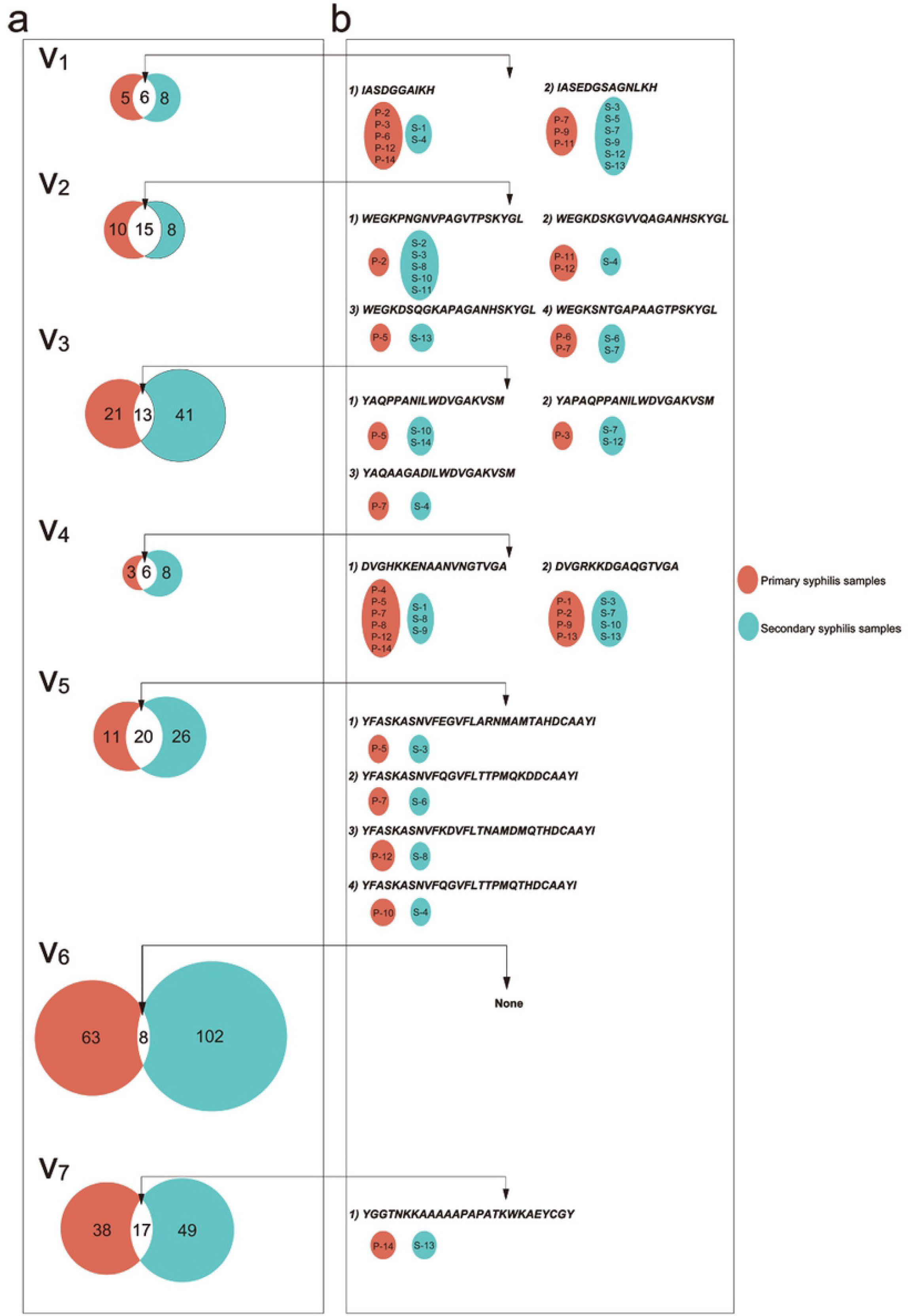
The scenario of same *tprK* amino acid sequences across the two-stage populations within the variable region of *tprK*. **a**, Each Venn diagram represents the ratio of same types of *tprK* amino acid sequence to all the exclusive types of amino acid sequences within the regions from their respective clinical samples. Numbers in each circle refer to the number of exclusive amino acid sequence types in primary and secondary syphilis samples. **b**, Graph displays the same types of amino acid sequences with a high proportion within each strain across the two-stage samples.

All seven V regions showed different degrees of variability between the two-stage populations, with a maximum obtained for the V6 region, in which we only detected up to seven identical types of amino acid sequences that continually appeared in the two-stage samples (Fig 4a). The ratios of the identical sequence types to the total exclusive sequence types obtained in primary and secondary syphilis samples were 12.7% (8/71) and 7.8% (8/110), respectively. Additionally, there was no case found where the predominant sequences were the same in V6 (Fig 4b). Then, the levels of nucleotide diversity in V6 between each sample (Dxy) were calculated using DnaSP v.6.12.01. As shown in Fig 5, the Dxy nucleotide diversity in V6 between each sample was almost above 0.15. This is in agreement with the proposed high diversity in V6 among most *T. pallidum* strains.

**Figure 5.**
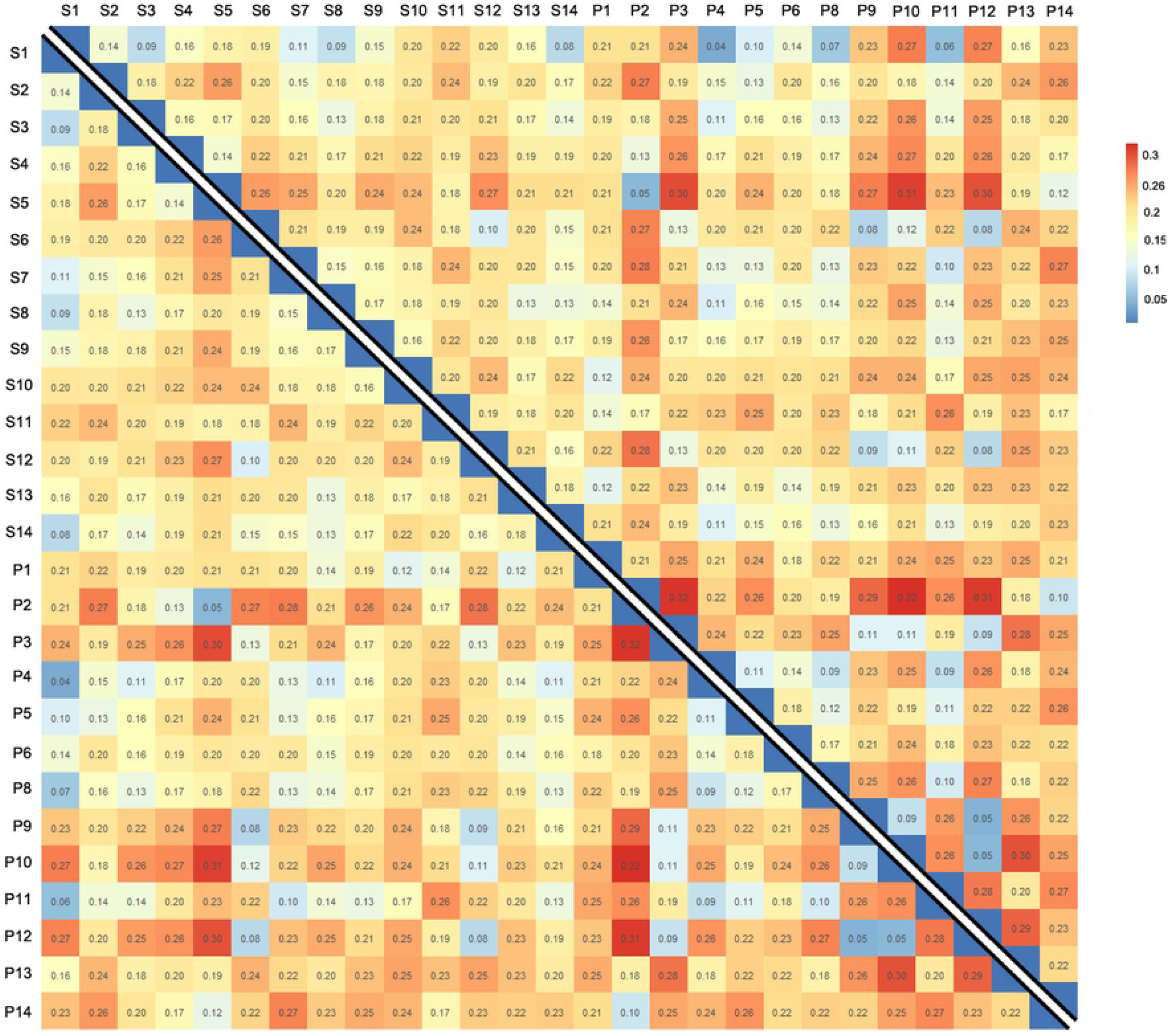
The diversity of nucleotide sequences in V6 at the inter-strain level. It was only one nucleotide sequence captured in P7 strain, so there were no sequence diversity data of P7 strain.

## Discussion

As a *tpr* gene family was identified in the Nichols strain of *T. pallidum* [19], the antigen-coding *tprK* has been extensively studied because of its highly variable antigenic profile [9, 10, 14, 20–22]. Similar to known mechanisms where many pathogens undergo antigenic variation to evade the immune system and establish chronic infection in the host [5, 23], *tprK* was believed to play an essential role in syphilis pathogenesis [15, 16]. Hence, efforts to understand *tprK* diversity in the context of human infection, particularly in different clinical stage samples, would be beneficial to clinically elucidate the role of *tprK* in successive episodes of this chronic infection and would contribute to a deeper understanding of the pathogenesis of syphilis.

In this study, by using NGS in combination with an in-house Perl script, we retrieved the variants within the seven regions of each strain and calculated the relative frequency of the variants, thus discovering that *tprK* always presented a predominant sequence and numerous minor variants within each V region in the two-stage samples. Moreover, most variants were found to be distributed in the 1-5% range, indicating that the diversity of *tprK* gene always consists of numerous minor variants that are beyond the detection limit of Sanger sequencing [24]. Interestingly, the variants with the frequencies of 20-80% within the variable regions of *tprK* were rare in primary syphilis samples; while the variants in this range were more frequently detected (except V1) in secondary syphilis samples, which was attributed to that the frequencies of predominant sequences within the V regions decreased and the minor variants in 10-60% range increased. This might imply that the dominant sequence of *tprK* in early infection would no longer allow *T. pallidum* to develop into the next stage. Then, the frequency of the predominant sequence in the variable regions (except V1) decreases, and a certain variant among numerous low-frequency variants is selected to increase its frequency in the populations [25]. Consequently, a new TprK is generated to evade the original antibody response, contributing to *T. pallidum’s* success of persistence in the host [16, 22]. Furthermore, probably owing to a strict 3 bp changing pattern in each variable region, there were no frame shifts in the *tprK* gene found in the two-stage samples [9, 18], suggesting that despite the high recombination rate occurring in the *tprK* gene of *T. pallidum*, an intact *tprK* ORF is always maintained, demonstrating that a normal TprK antigen is essential for *T. pallidum* to make a living as a stealth pathogen.

Although more distinct nucleotide sequences were found within each variable region of *tprK* in secondary syphilis patients, most were found in the same region, V6, and the least amount of sequences was found in V1. A remarkable inter-population redundancy was observed in the amino acid sequence level of *tprK* among the two-stage samples, which is similar to previous studies [9, 26]. This finding might indicate that *tprK* has superior sequences that are better for *T. pallidum* to infect the host throughout evolution in different patients. Among all V regions, V1 had a strong inter-population sharing ability; moreover, some amino acid sequence types in V1 continually appeared in the two-stage samples. Notably, the amino acid sequences IASDGGAIKH and IASEDGSAGNLKH were not only the ones that have a high proportion in inter-population sharing but also the ones that always had a relatively high frequency (above 80%) in the populations. In fact, the sequence IASDGGAIKH was found to be the most stable amino acid sequence of inter-population redundancy in Pinto’s study [26]. It was shared by seven strains, five directly from primary syphilis samples and two from the primary to the secondary stage samples. The sequence IASEDGSAGNLKH also showed a high sharing capacity, although it was found only in primary syphilis samples in the study [26]. Similar findings suggest that V1 would be a promising region for researchers to focus on. A previous study [27] found that the molecular localization in the N-terminal region (containing the region V1) of *tprK* displays promising partial protection in a rabbit model. Therefore, the most stable peptides in V1 could be a potential vaccine component.

In this study, we also found that V6 always presented high heterogeneity at the intra-strain level in contrast to other regions. The existence of a highly diverse region in this antigen-code gene of an isolate of *T. pallidum* may greatly enable the pathogen to resist binding by existing opsonic antibodies and less likely to be recognized by activated macrophages [16]. The amino acid sequences of V6 also showed high diversity at the inter-strain level, which seemed to represent an isolate-specific pattern. This could probably explain why the protection of TprK was compromised and the lack of heterologous protection [22]. Additionally, it can be very difficult to distinguish between treatment failure (relapse) and reinfection in clinical practice. Whether there is speculation that the sequences in V6 retain a high degree of similarity in a relapse case, but the sequences are highly isolate-specific in a reinfection case? This speculation requires further supporting evidence from additional experiments in the future.

Finally, the limitations of our research should be discussed. First, the study did not explore the function of sequences within each V region, and it remains to be explored in the future. Second, this study lacked the direct observation of the entire process of variations of *tprK* within disease development. A rabbit model might be needed to monitor the dynamic change in *tprK*. Third, the study provided information on individual V regions instead of information on a single *tprK* ORF. This could result in artificial sequences, if it is assumed that certain nucleotide sequences within the V regions are derived from a same single *tprK* ORF.

In this study, we revealed that the characteristic profile of *tprK* was different in primary and secondary syphilis patients, demonstrating that throughout the development of the disease, *T. pallidum* constantly varies its *tprK* gene to obtain the best adaptation to the host. Interestingly, *tprK* always maintains a feature in these ongoing variations, that is, having a relatively conserved region (V1) and a highly diverse region (V6). These findings could provide important information for unveiling the mysterious role of *tprK* in persistent syphilis infection and for further exploring the promising potential vaccine components.

## Materials and Methods

### Ethics statement

The subjects of the study were adults, and all provided written informed consent in accordance with institutional guidelines prior to the study. This study was approved by the Institutional Ethics Committee of Zhongshan Hospital, Medical College of Xiamen University, and it was in compliance with national legislation and the Declaration of Helsinki guidelines.

### Collection of clinical materials

Ulcer swab samples were collected from patients with primary syphilis (P1~14) (the samples included in this study were the same as the samples used previously [18]) and secondary syphilis (S1~14). The clinical diagnosis of syphilis was based on the US Centers for Disease Control and Prevention (CDC) [28] and the European CDC (ECDC) guidelines [29]. The clinical data, including patient age, gender, syphilis serology results and clinical manifestations, were also collected at Zhongshan Hospital, Xiamen University.

### Isolation of DNA and detection of treponemal DNA

DNA was extracted from swab exudates using the QIAamp DNA Mini Kit (Qiagen, Inc., Valencia, CA, USA), according to the manufacturer’s instructions, and careful precautions were implemented to avoid DNA cross-contamination between samples[30]. Each sample was quantified by targeting *tp0574* through qPCR using a 96-well reaction plate with a ViiA 7 Real-Time PCR System (Applied Biosystems, USA) according to the protocals already established in the laboratory [31]. The DNA samples, which were positive after qPCR targeting of *tp0574*, were subjected to amplification of *tp0136* to determine whether these 28 clinical strains belonged to the Nichols-like group or SS14-like group [32].

### Segmented amplification of the *tprK* gene

The segmented amplification of the *tprK* gene was conducted as previously described[18], with the extracted DNA being directly used for the amplification of the *tprK* full open reading frame (ORF) and the amplicons being gel purified. The purified full-length *tprK* amplicons was used as a segmented amplification template for partial amplification of four fragments of 400-500 bp (overlapping by at least 20 bp), covering the *tprK* ORF. The size of all the products was verified by 2% agarose gel electrophoresis, and the products were gel purified. A high-fidelity PCR polymerase, KOD FX Neo polymerase (Toyobo, Osaka, Japan), was used for amplification and the amplification primers are list in S5 Table.

### Library construction and next-generation sequencing

The four subfragment amplicons corresponding to each sample were mixed in equimolar amounts into one pool to produce a separate library, using a barcode to distinguish each sample. Library construction and sequencing were performed by the Sangon Biotech Company (Shanghai, China) on the MiSeq platform (Illumina, San Diego, CA, USA) in paired-end bi-directional sequencing (2×300 bp) mode. The FastQC (http://www.bioinformatics.babraham.ac.uk/project/fatsqc/) and FASTX (http://hannonlab.cshl.edy/fastx_toolkit) tools were applied to check and improve the quality of the raw sequence data, respectively. The final reads of *tprK* collected from 28 isolates were compared with the *tprK* of the Seattle Nichols strain (GenBank accession number AF194369.1) using Bowtie 2 (version 2.1.0).

Based on the previously described principle that was used to extract sequence data [18, 26], an in-house Perl script was developed and applied to specifically capture DNA sequences within seven regions of the *tprK* gene across 28 strains from raw data, both forward and reverse. Thus, the exact number of distinct sequences within the seven variable regions of the *tprK* gene from each sample was acquired. The intrastrain heterogeneous sequences were valid if the following conditions were simultaneously verified for any variant sequence: 1) being supported by at least fifty reads and 2) displaying a frequency above 1%. Then, the relative frequency of the sequences within each variable region was calculated.

### Accession numbers

The raw data sequences of *tprK* in this study were deposited in the SRA database (BioProject ID: PRJNA498982 and PRJNA512914) under following BioSample accession numbers: SAMN10340238-SAMN10340251 for P1~14 and SAMN10690826-SAMN10690839 for S1~14, respectively.

## Acknowledgments

We thank the patients all for their support in this research and thank the Zhongshan Hospital colleague for their assistance in collection of research data.

## Supporting information

**S1 Table. The characteristics of clinical samples and the background data of *tprK* by NGS**

**S2 Table. The nucleotide sequences within the seven variable regions (V1-V7) of *tprK* captured directly from 28 clinical samples** (A) Captured nucleotide sequences ranked according to their relative frequency within each population from 14 primary syphilis samples (P1~14) and (B) from 14 secondary syphilis samples (S1~14)

**S3 Table. The amino acid sequences within the seven variable regions (V1-V7) of *tprK* captured directly from 28 clinical samples** (A) The amino acid sequences were from 14 primary syphilis samples (P1~14) and (B) the amino acid sequences were from 14 secondary syphilis samples (S1~14). * indicates synonymous nucleotide sequences within the same strain

**S4 Table. The scenario of same *tprK* amino acid sequences across the two-stage populations within the variable regions (V1~V7)**

**S5 Table. The primers for *tprK* amplification**

